# No evidence for unknown archaic ancestry in South Asia

**DOI:** 10.1101/068155

**Authors:** Pontus Skoglund, Swapan Mallick, Nick Patterson, David Reich

**Affiliations:** Department of Genetics, Harvard Medical School, Boston, MA, USA; Broad Institute of Harvard and MIT, Cambridge, MA, USA; Howard Hughes Medical Institute, Boston, MA, USA

## Abstract

Genomic studies have documented a contribution of archaic Neanderthals and Denisovans to non-Africans^1,2^. Recently Mondal *et al.*^3^ published a major dataset—the largest whole genome sequencing study of diverse South Asians to date—including 60 mainland groups and 10 indigenous Andamanese. They reported analyses claiming that nearly all South Asians harbor ancestry from an unknown archaic human population that is neither Neanderthal nor Denisovan. However, the statistics cited in support of this conclusion do not replicate in other data sets, and in fact contradict the conclusion.

The main evidence cited by Mondal *et al.* is statistics suggesting that indigenous Andamanese and mainland Indian groups share fewer alleles with sub-Saharan Africans than they do with Europeans and East Asians; such statistics have previously been reported in Australasians, for whom they represent key evidence of Denisovan admixture^2^. To document their signal, Mondal *et al.* compute *D*-statistics^1,4^ of the form:

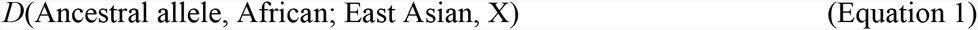

These statistics test the hypothesis of an equal rate of allele sharing of East Asians and X with Africans. In Mondal *et al.*’s computation, these statistics are negative when X is any Indian group or Andamanese, a result they interpret as evidence of more archaic ancestry than in East Asians. As they find no evidence of excess allele sharing with Neanderthals or Denisovans, they argue that the contribution is from an unsampled archaic lineage.

We sought to replicate these statistics in two single nucleotide polymorphism (SNP) data sets of ∼600 thousand SNPs each^5,6,4,7^, whole genome sequence data from the 1000 Genomes Project phase 3 (∼78 million SNPs)^8^, and high-coverage genomes (∼34 million SNPs)^9^. There is no evidence for excess archaic ancestry in South Asians in any of these four data sets (Figure 1), and in fact the values reported by Mondal *et al.* (*D*=-0.024 ± 0.004; Supplementary Table 13 of their study^3^) are inconsistent with those in each of these other datasets (all P < 10^-5^ by a one-tailed test).

**Figure 1.**
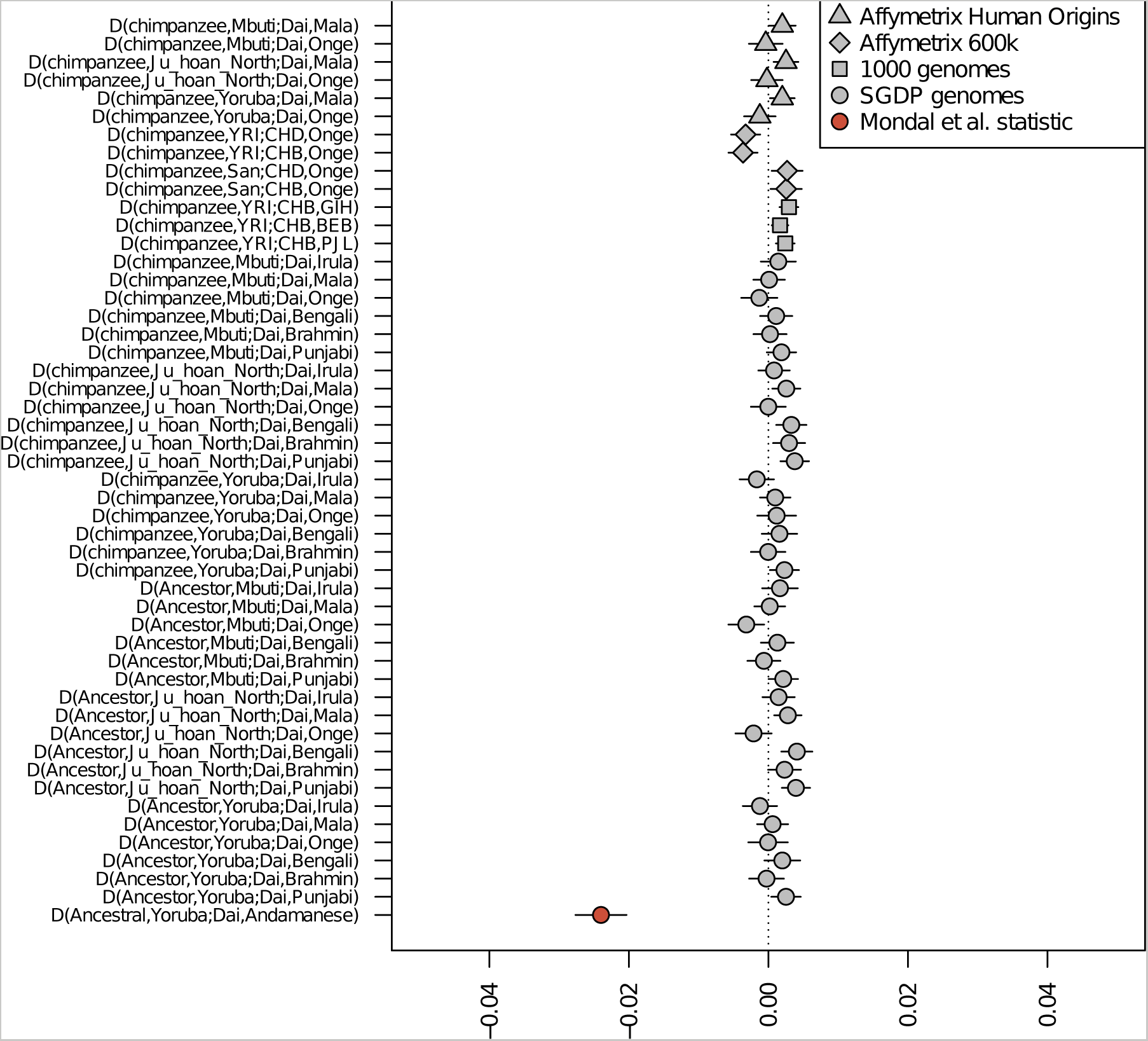
The key statistic used to support the claim of unknown archaic ancestry in Andamanese and mainland Indians by Mondal *et al.* is inconsistent with all previously published datasets. Evidence for unknown archaic ancestry in Andamanese and mainland Indians does not replicate in four previously published data sets. Error bars show 1 standard error on each side. All statistics except the one reported by Mondal *et al*. are consistent with no excess archaic admixture in Indians (│Z│ < 2).

Mondal *et al.* also report statistics suggesting more archaic ancestry in indigenous Australians than in indigenous Papuans, as reflected in *D*-statistics that are far more skewed from zero when X=Australian than when X=Papuan^3^. They interpret this as evidence of unknown archaic ancestry in Australians to a greater extent than in Papuans. However, we do not replicate this excess when recomputing this statistic using high-coverage genomes from these populations: *D*(chimpanzee, Yoruba; Dai, Australian) = -0.031 ± 0.003 and *D*(chimpanzee, Yoruba; Dai, Papuan) = -0.029 ± 0.003. In addition, a direct comparison between Australians and Papuans provides no evidence for a difference: *D*(chimpanzee, Yoruba; Australian, Papuan) is only Z=0.6 standard errors from zero^9,11^. These findings support the notion that Papuans and Australians descend from a homogeneous ancestral population, and are inconsistent with the suggestion that Australians harbor much more archaic ancestry than Papuans.

In fact, some of the statistics computed by Mondal *et al.* directly contradict their proposed model of unknown archaic ancestry specific to Indians and Andamanese. Figure 1b of Mondal *et al.* suggest that the Riang (RIA)—a Tibeto-Burman speaking group from the northeast of India for which sequencing data are newly reported in the study—derive almost all of their ancestry from the same East Asian lineages as populations like Dai and Han Chinese, which in Figure 1b of Mondal *et al.* have no evidence of unknown archaic ancestry. Under the authors’ hypothesis of more archaic ancestry in lineages that are unique to South Asia than in lineages shared with East Asians, one would not expect a significant statistic in the Riang, but in fact the signal is just as strong as it is for the Andamanese Onge, Andamanese Jarawa, mainland Irula and mainland Birhor, the great majority of whose ancestry is inferred to derive from lineages unique to South Asia.

One possible explanation for the skew that the authors observe^3^ is batch artifacts, reflecting differences in laboratory or computer processing between the data newly reported by Mondal *et al.,* and the data from non-Indians used for comparison^10^. Separate processing is known to be able to cause correlation of errors within datasets, and this could explain why the newly reported South Asian genomes appear (artifactually) to share fewer alleles with other modern humans. However, the data used by Mondal *et al.* have not been made available for independent reanalysis, and without this, a definitive explanation is not possible. Whatever the explanation, our analyses contradict the claim of unknown archaic ancestry in South Asians.

## Acknowledgements

P.S. was supported by the Swedish Research Council (VR grant 2014-453). N.P. and D.R. were supported by NIH grant GM100233 and D.R. is a Howard Hughes Medical Institute investigator.

